# Use of Nanosphere Self-Assembly to Pattern Nanoporous Membranes for the Study of Extracellular Vesicles

**DOI:** 10.1101/758391

**Authors:** Marcela Mireles, Cody W. Soule, Mehdi Dehghani, Thomas R. Gaborski

## Abstract

Nanoscale biocomponents naturally released by cells, such as extracellular vesicles (EVs), have recently gained interest due to their therapeutic and diagnostic potential. Membrane based isolation and co-culture systems have been utilized in an effort to study EVs and their effects. Nevertheless, improved platforms for the study of small EVs are still needed. Suitable membranes, for isolation and co-culture systems, require pore sizes to reach into the nanoscale. These pore sizes cannot be achieved through traditional lithographic techniques and conventional thick nanoporous membranes commonly exhibit low permeability. Here we utilized nanospheres, similar in size and shape to the targeted small EVs, as patterning features for the fabrication of freestanding SiN membranes (120 nm thick) released in minutes through a sacrificial ZnO layer. We evaluated the feasibility of separating subpopulation of EVs based on size using these membranes. The membrane used here showed an effective size cut-off of 300 nm with the majority of the EVs ≤200 nm. This work provides a convenient platform with great potential for studying subpopulations of EVs.

## 1 Introduction

Cells are known to secrete extracellular vesicles (EVs) ranging in size from 30 to 1000 nm.^1^ These EVs include apoptotic, and microvesicles as well as exosomes. Apoptotic bodies (50 – 1000 nm) are released during cell death containing portions of the fractionated cell.^2^ Microvesicles (100 – 1000 nm) originate from plasma membrane budding while exosomes (30 – 150 nm) are produced intracellularly inside multivesicular endosomes or multivesicular bodies.^1–4^ Exosomes contain proteins and nucleic acids and take part in cellular communication, these small EVs can be found in bodily fluids and cell culture media. They carry a fingerprint from their originating tissue creating a valuable opportunity for disease understanding and diagnosis, particularly of cancer.^3–14^ Moreover, they could potentially be used in therapeutics as healthy cells shed exosomes as well.^15–18^

The lack of a platform that offers the ability to compare side-by-side different subpopulations of EVs, in isolation and cellular co-culture systems, continues to stall our overall knowledge of EVs and their effects on cellular communication.^1,19^ The preferred methodology for the isolation of small EVs is ultracentrifugation although many others continue to emerge such as membrane based isolation.^4,20–29^ Additionally, the most commonly used methodology for the study of small EVs in cellular co-culture is the use of commercially available track etched membranes. These membranes are thick rendering them low in permeability and often contain merged pores.^30–33^

A membrane-based approach is valuable since it represents a dual purpose platform which can be used in isolation and cellular co-culture systems for the study of small EVs. The thickness of these membranes is not trivial given the fact that in filtration devices designed for nanoscale species the small pores inherently increase the fluid resistance across the membrane.^34^ High permeability, at the nanoscale, can be obtained through membranes with a pore size to thickness ratio close to one.^34,35^ Such platform should provide ultrathin membranes with close control on pore size over the range of small EVs. Another reason for minimal thickness is to improve tissue barrier function in a direct co-culture format by allowing physical contact and fast exchange of soluble factors,^36–38^ establishing a more physiologically relevant model. Several publications have reported on processes for the production of nanoscale pores but often produce thick membranes, do not successfully demonstrate membrane release, or are based on patterning approaches that do not offer control over the full range of EVs sizes.^35,39–51^ Publications reporting on the fabrication of ultrathin membranes with control over pore size in the range of EVs heavily rely on microfabrication technologies requiring harsh chemicals and time consuming steps.^52,53^

Our research group has demonstrated the fabrication of large area ultrathin nanoporous silicon nitride and microporous silicon oxide membranes obtained by solid phase crystallization and traditional patterning methods, respectively. We have also demonstrated the viability of utilizing those membranes for cell culture studies. ^38,54^ In the current work, we utilize a patterning approach with close control of pore size over the range of small EVs, together with a simple methodology for the release and integration of the ultrathin membrane into a silicone based device for the study of EVs. To the best of our knowledge, such a comprehensive process has not yet been demonstrated. This platform will impact cellular co-culture studies and isolation based methodologies including the previous research published by our group on the capture and release of EVs in a tangential flow microfluidic device.^55^

Here, we report the fabrication of ultrathin free-standing nanoporous membranes. These membranes can be made of SiN for increased mechanical robustness or SiO_2_ for optimum optical transparency and surface functionalization.^38,54^ The pores have been patterned through the use of self-assembled nanospheres, similar in size and shape to small EVs. This patterning technique is a bottom-up approach known as nanosphere lithography (NSL) and offers pore size control over a large area at an affordable cost.^35^ The most commonly used techniques for nanosphere self-assembly are spin coating and interfacial trapping. The latter approach allows for efficient use of the nanospheres, with virtually no waste, while spin coating requires a large volume most of which is ultimately lost. Interfacial trapping can render close packed or non-close packed arrangements when using neutral or charged nanospheres, respectively. A non-close packed arrangement allows for the independent control of pore size and porosity.^51,56^ We adopted the latter methodology because of its versatility and reproducibility.

The last step in membrane fabrication is the release or lift-off that detaches the membrane from its support, often a silicon wafer, rendering it free-standing. This process usually requires strong chemicals and is time consuming. Reducing the cost can be achieved by using thinner silicon wafers, however these are more prone to breakage. The use of thin films acting as sacrificial layers, which can be dry or wet etched, has been explored mainly in the electronics industry but also for membrane release.^54,57–60^ We adopted the use of a zinc oxide (ZnO) thin film as a sacrificial layer due to its low-cost, stability through the fabrication process, and fast etching rate in mild chemistry. We have integrated our membrane into a simple silicone device and evaluated their size cut-off characteristics and permeability of EVs. These membranes are useful for the isolation and recovery of EVs for their analysis as well as for the modulation of EV-mediated communication in co-culture. We expect this platform to aid researchers in contributing to the body of knowledge surrounding the therapeutic and diagnostic potential of the different subtypes of EVs.

## 2 Experimental details

### 2.1 Membrane fabrication

The process followed for membrane fabrication has been depicted in Figure 1 and will be described in detail in the following sub-sections. The versatility of this methodology enables it to adapt to varied specific needs. The initial size of the nanospheres can be easily changed as well as their composition which allows the use of alternative chemistries for size reduction and control over a different pore size range. The membrane composition can be substituted by a myriad of materials as long as selectivity under HCl is maintained. The material used as overcoat to form the nanoporous etching mask can also be substituted for other metals and even oxides to modify the porous pattern transfer or the composition at the surface. Finally, it is worth noting that there is no inherent limitation to scale the process to smaller or larger samples.

**Figure 1.**
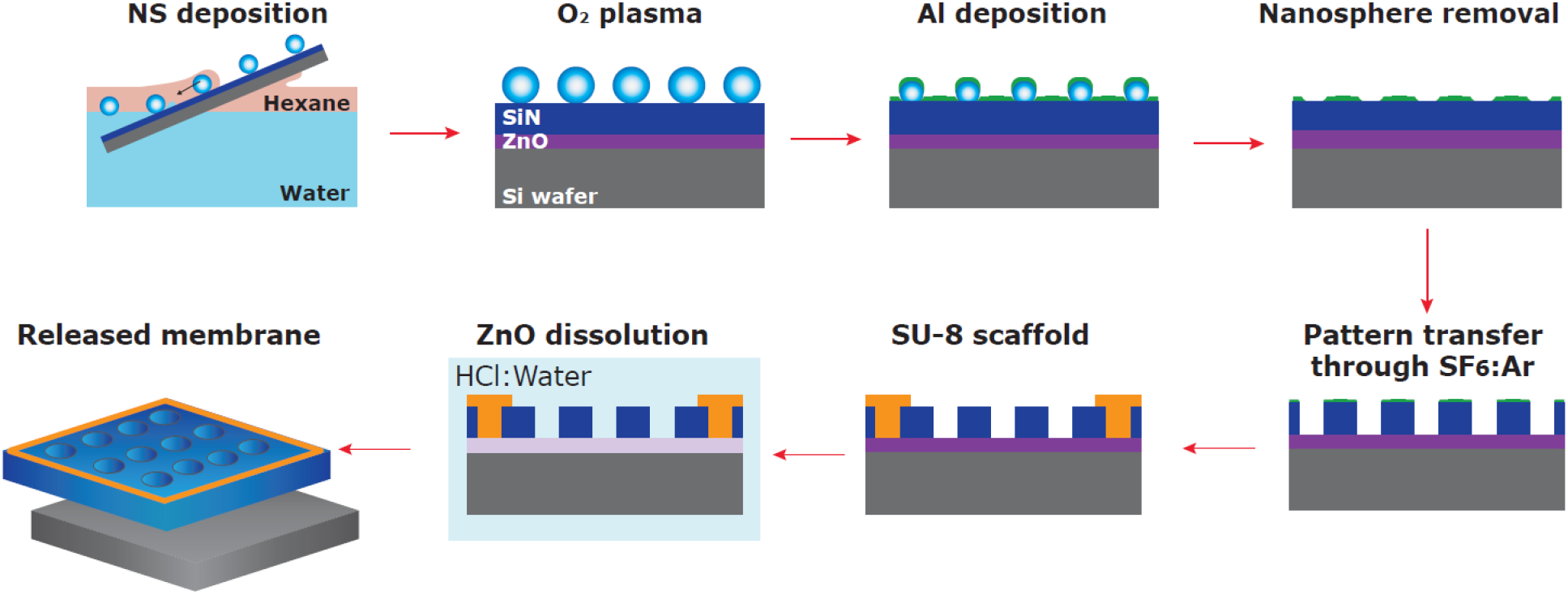
Ultrathin nanoporous membrane fabrication process. Schematic summary of the integrated process, from nanosphere self-assembly and transfer to membrane release.

**Figure 2.**
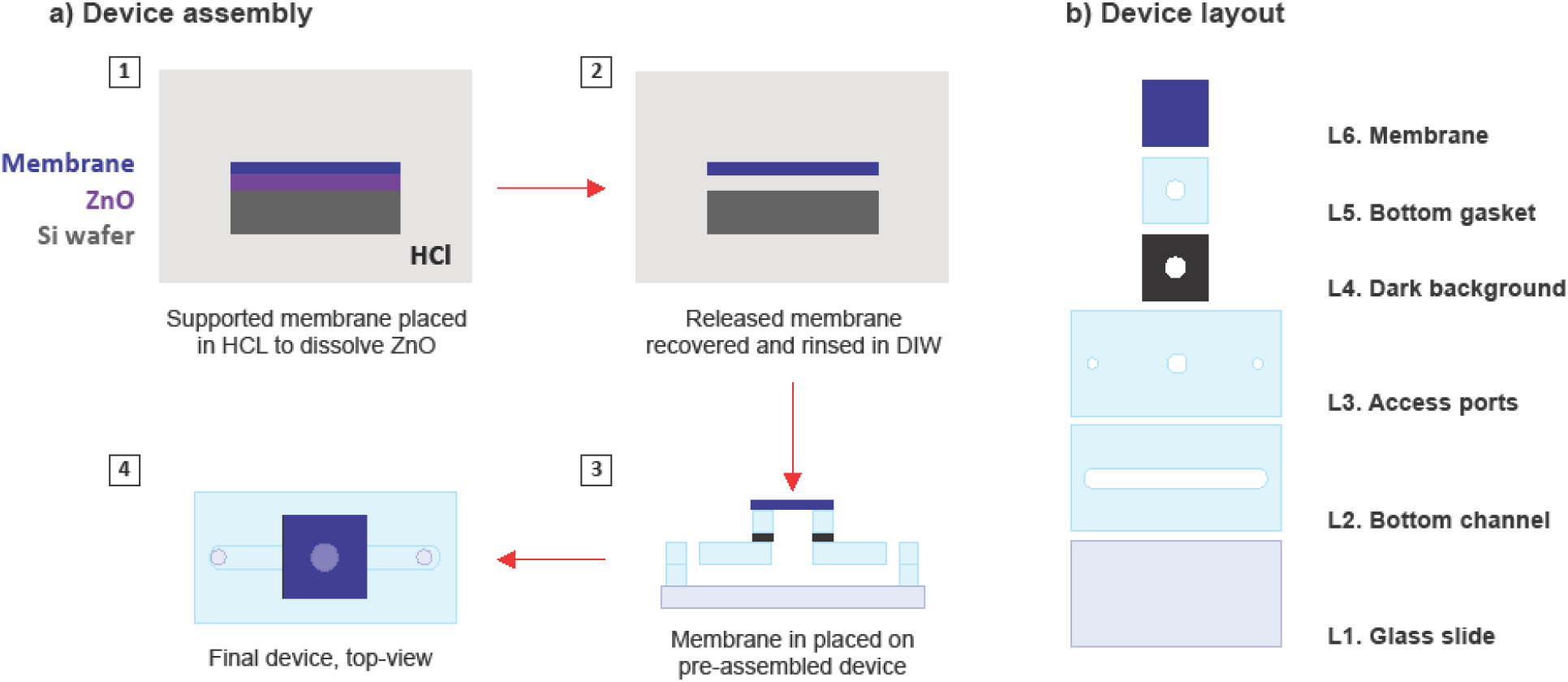
Membrane integration: a) overall device layout, and b) schematic representation of the process flow followed to release the supported membrane into a silicone device.

#### Substrate preparation

All substrates used were silicon wafers with <100> orientation purchased from University Wafer (Massachusetts, United States) coated with 100 nm of ZnO deposited at room temperature from a Zn target purchased from Kurt J. Lesker (Pennsylvania, United States) by reactive sputtering (Kurt J. Lesker, PVD-75) in an Ar:40%O_2_ atmosphere, this ZnO film was annealed for 1 h at 600 °C under N_2_ flow in a Bruce furnace. The ZnO film is used as a sacrificial layer for membrane release. The substrate were then coated with 120 nm of low-stress silicon nitride deposited by plasma enhanced chemical vapor deposition (Rogue Valley Microdevices – Oregon, United States).

#### Nanosphere self-assembly and deposition

Positively charged polystyrene nanospheres were purchased from Invitrogen (California, United States) with a nominal size of 300 nm in diameter. The aqueous stock solution was diluted in a mixture of isopropanol and ultrapure water (4:6 v:v). The substrate was mounted on a holder with a 10° upward tilt and immersed in a polypropylene beaker containing 175 ml of ultrapure water. Next, 25 ml of n-hexane were carefully added through the sidewall of the beaker to form an oil/water interface. Then, 200 μl of the diluted stock solution were injected into the hexane phase, 1 mm above the interface, at a speed of 50 μl/min controlled with a syringe pump. The system was then covered with a lid and the nanoshperes were allowed to self-assemble for 12 mins. Finally, the substrate was withdrawn at a controlled speed in the range from 25 to 450 μm/s. The withdrawal speed was controlled by attaching the substrate holder to a syringe pump. The NS attach to the substrate as it travels across the n-hexane/water interface. The methodology described for nanosphere self-assembly has been adapted from previous publications.^51,61,62^ Here, the self-assembly and transfer were carried out inside an enclosure maintained around 50 % relative humidity, this parameter was found to be crucial in ensuring repeatability. After the nanosphere monolayer was deposited on the substrate, the samples were allowed to completely dry overnight at room temperature.

#### Nanoporous etching mask fabrication

The NS monolayers were exposed to an oxygen plasma for size reduction. Because the size of the NS determines the final size of the pore, this plasma treatment allows for pore size control. The samples were placed in a reactive ion etch (RIE) chamber from South Bay Technology (California, United States) which was filled with O_2_ to a pressure of 130 mTorr (≈ 45 sccm O_2_) and supplied with a power of 130 W. The etch rate under these conditions was found to follow a linear trend with an approximate slope of 6 nm/s. After NS size reduction, an Al thin overcoat was deposited by thermal evaporation in a Kurt J Lesker tool. Scotch tape was used to mechanically remove most of the nanospheres and then the samples were sequentially sonicated in toluene and acetone for 10 mins, twice in each solvent. Finally the samples were cleaned in isopropyl alcohol followed by ultrapure water and dried under an air stream. After nanosphere removal, the Al thin film exhibited a nanoporous pattern and was used as etching mask.

#### Nanoporous pattern transfer

The pattern was transferred into the SiN layer through dry etching. The samples were introduced into the RIE chamber filled with Ar:SF_6_ (1:2 sccm) to a pressure of 120 mTorr and a supplied power of 175 W. The etch rate under these conditions was found to be ≈ 1 nm/s. Next, the Al was fully removed by immersing the samples in a 1 M solution of NaOH for 1 min.

#### Polymeric scaffold

The scaffold material is SU-8 (3025 from MicroChem Corp. Massachusetts, United States), a commonly used negative resist. The samples were primed by immersing them in 1M HCl for 5 secs which etches the ZnO exposed at the bottom of the pores and significantly increases scaffold attachment by improving mechanical gripping. The SU-8 was deposited by spin coating through a two-step process. First, spinning at 500 rpm for 10 s with an acceleration of 100 rpm/s. Then, spinning at 3000 rpm for 45 s with an acceleration ramp of 100 rmp/s. Following SU-8 deposition, a soft bake was performed for 10 mins at 95 °C. Then the samples were exposed with a dose of 225 mJ/cm^2^ in a Karl Suss 1X mask aligner equipped with a broadband lamp. Next, a post exposure bake was performed; first at 65 °C for 1 min and then at 95 °C for 3 mins with a waiting period of 10 mins between these steps. Finally, the samples were developed under mild agitation for 5 mins and rinsed with isopropanol. All heating steps were performed with a ramp of 5 °C/min from a starting temperature of 60 °C, this minimizes mechanical strain that can compromise SU-8 attachment. The scaffold exhibits a grid pattern which is 20 μm thick with 100×100 μm openings and 10 μm wide struts.

We have also used the methodology described above for the fabrication of SiO_2_ membranes. The specific details of the SiO_2_ deposition and post-deposition treatment are described in the supporting information which also includes electron micrographs of the porous membranes **(Figure S1**, **Supporting information)**.

### 2.2 Device integration

The silicone based device consisted of a bottom channel, a membrane chip, and a top chamber. The device was fabricated from 600 μm thick silicone sheets purchased from Silicone Specialty Fabricators (California, United States) custom cut with the use of a Silhouette Cameo digital craft cutter. The bottom channel (2×17 mm) was plasma bonded to a clean glass slide followed by another layer with two access ports (2 mm diameter) and one central opening (3 mm diameter) to create an interaction area connecting top and bottom chambers. A silicone gasket with a central opening (3 mm diameter) with a backing made from black vinyl was secured to the bottom channel central opening with the use of Kapton tape. This silicone gasket was the layer receiving the membrane after lift-off. The supported membrane was immersed in 1 M HCl for lift-off. It is important to note that this lift-off process only takes a couple of minutes, which is a remarkable improvement over the through wafer etching commonly used. After release, the free-standing membrane was carefully removed with tweezers and rinsed in deionized water **(Video S1, Supporting information)**. The free-standing membrane was then placed over the plasma treated silicone gasket and allowed to bond at room temperature for 2 hrs. The top chamber consisted of a silicone piece with a 4 mm center opening that was plasma treated and bonded to the membrane. Finally, silicone tubing with an internal diameter of 0.031” (Cole-Parmer - Illinois, United Stated) was affixed to the bottom channel outlets.

### 2.3 Size cut-off evaluation

The fabricated device operates under convective flow by creating a simple siphon to pull the fluid from the top chamber, through the membrane, into the bottom channel, and out for collection. The device was set onto the stage of a fluorescent microscope (Leica Microsystems DMI6000, Wetzlar, Germany), then the top chamber and bottom channel were filled with ultrapure water, including the silicone tubing sections which serve as bottom channel inlet and outlet. To initiate flow a 30 cm height different was set between the fluid level inside the top chamber and the bottom channel outlet, while constricting the flow through the bottom channel inlet. After most of the ultrapure water was depleted, the top chamber was refilled with a testing solution containing either 200 or 300 nm fluorescent particles (unmixed). The testing solution was allowed to permeate through the membrane and into the bottom channel where the fluorescent intensity was tracked as a function of time. The obtained values were adjusted to the background intensity and normalized to the intensity at the membrane, the data has not been adjusted for quenching effects. The fluorescent latex beads were purchased from Magsphere Inc (California, United States) with nominal particle sizes of 200 nm (green fluorescent) and 300 nm (red fluorescent).

### 2.4 Filtration of extracellular vesicles

EVs were purchased lyophilized from HansaBioMed (Tallinn, Estonia) and reconstituted according to their specifications. Next, we fluorescently labelled the EVs with CFSE dye (5(6)-carboxyfluorescein N-hydroxysuccinimidyl ester) purchased from Invitrogen (California, United States).^63^ This dye, commonly used for the tracking of cells as well as EVs, permeates through the membrane and interacts with free amine to generate fluorescent protein conjugates which are much less membrane permeable.^63–69^ Moreover, we utilized the CFSE dye because it has been shown to preserve the size characteristics of the EVs.^70^ In order to prevent non-specific adsorption due to charge-based interaction between the membrane and the EVs, we coated the membrane with bovine serum albumin (BSA). A 5 mg/ml BSA solution prepared in phosphate buffered saline (PBS) was clarified at 2000 rpm for 8 mins and loaded onto the top and bottom chambers of the device. After 1 hr, the device was thoroughly rinsed with clean PBS. Similar BSA treatment has been previously shown to yield a 3.5 nm thick coating,^40^ this represents a 3% pore size decrease in our membrane which does not represent a significant change. The labelled EVs were loaded onto the top chamber and allowed to permeate through in the same fashion as the experiments described in the subsection 2.3. After the experiment, the filtered fraction was recovered for further characterization by nanotracking analysis (NTA).

### 2.5 Imaging

Scanning electron micrographs were taken with a Tescan Mira3 (Brno, Czech Republic) or an S-4000 Hitachi Ltd (Tokyo, Japan) both equipped with field emission electron guns. The samples were coated with ~ 10 nm of sputtered gold to reduce charging effects.

## 3 Results and discussion

### 3.1 Withdrawal speed optimization

Many authors have described methods for successful monolayer transfer onto varied substrates.^56,71^ Although the forces involved are well understood, the transferring methods vary significantly. As the substrate travels through an interface, several parameters come into play to adequately balance attractive and repulsive forces.^56,71^

Charged nanospheres are irreversibly trapped at the water/hexane interface where they form dipoles caused by the surface functional groups dissociating. These dipoles will spontaneously align normal to the interface, locking the nanospheres and ensuring the presence of a monolayer (dipolar interaction). Furthermore, electrostatic repulsion forces the nanospheres to repel each other, allowing the formation of a non-close packed monolayer (Coulomb interaction). These repulsive forces are mainly responsible for stabilizing the nanospheres at the interface. Attractive capillary forces, however, play a crucial role during the transfer of the monolayer onto a substrate and should be avoided to prevent aggregation. This phenomena takes place twice during the process, first as the substrate is pulled through the interface, and second as the transferred monolayer dries undisturbed. Because the relative humidity directly affects the drying process, we first set this parameter constant at 50 % for all experiments. Aggregation during substrate withdrawal is the result of two interlocked parameters: speed and tilt of the substrate. Large tilt angles (more vertical) are usually avoided since they require nanosphere monolayer compression to maintain inter-nanosphere spacing. We optimized the withdrawal speed while keeping the substrate tilt (10°) constant. Optical and electron micrographs of the samples obtained at four different speeds (25, 250, 450, and 900 μm/s) are shown in Figure 3 for the 300 nm nanospheres used; we found similar results for 200 nm nanospheres **(Figure S2**, **Supporting Information)**. We found that slow speed (25 μm/s) promotes the formation of folds in the monolayer which are observed as stripes. These folds are most likely due to oscillations of the interface resulting in secondary deposition of nanospheres. These defects, aligned parallel to the interface, are long range defects that locally affect the inter-nanosphere spacing and result in aggregation within the fold. Faster speed (900 μm/s) causes the inter-nanosphere spacing to vary significantly along the sample. Moreover, fast speed increases the probability of trapping water resulting in localized aggregation. Due to minimal presence of large size defects and aggregation, a withdrawal speed of 250 μm/s was selected for further membrane fabrication. These optimized conditions can be reliably used with Si, SiO_2_, and SiN. It is worth noting that we only observed significant difference on nanosphere arrangement and transfer on surfaces with water contact angles <5°. These super hydrophilic surfaces, such as oxygen plasma treated SiO_2_ and parylene, do cause water to remain on the surface after withdrawal with the concomitant effect of nanosphere aggregation. Hydrophobic surfaces, such as HF cleaned Si, with a water contact angle >80° remain unaffected by this phenomena.

**Figure 3.**
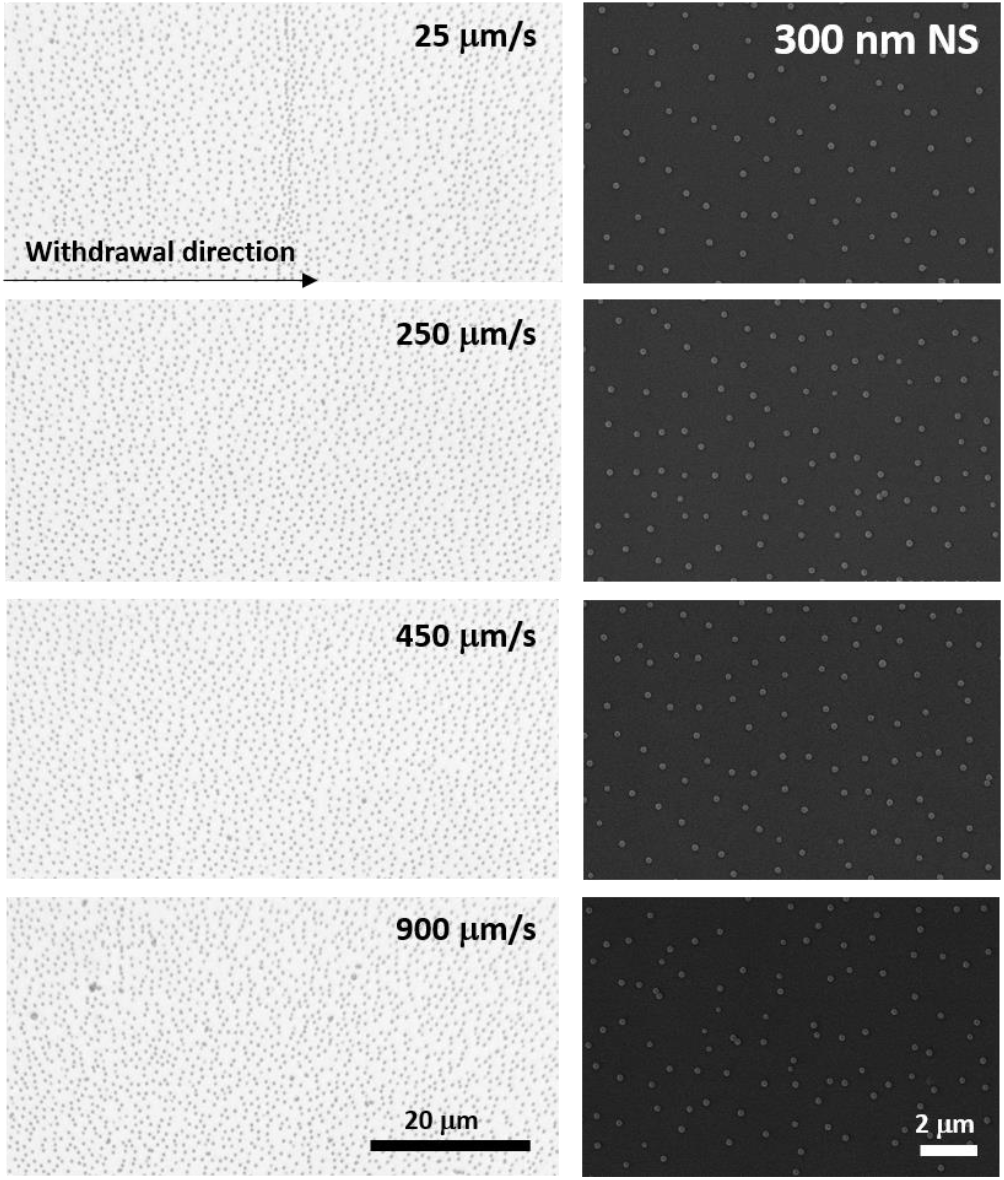
Withdrawal speed optimization. Optical and electron micrographs taken from 300 nm self-assembled monolayers transferred onto the substrate at different withdrawal speeds.

### 3.2 Size control

It is important to note that perfect self-assembly and pristine transfer onto a different substrate without defects is experimentally impossible. Instead, minimization of defects should be implemented through an application driven optimization parameter. In the context of membranes, the most important defects are the ones pertaining to pore size. Aggregates of nanospheres cause the average pore size to increase and the size dispersion to spread. Withdrawal speed optimization together with post-transfer treatment can greatly reduce the presence of aggregates. The use of oxygen plasma has been previously applied to reduce the size of polystyrene nanospheres.^72–74^ The dry etching profile is isotropic in nature and can be used to minimize defects such as aggregates. As shown in Figure 4a, we found the nanosphere diameter to linearly decrease in the etching range tested (0 – 30 s), with an approximate reduction rate of 7 nm/s. Figure 4b shows a top-view of the reduced NS, the etch is anisotropic with the height of the nanospheres decreasing faster than the width resulting in nanospheres than flatten over time. This effect can be seen in the tilted SEM micrographs included in the supporting info for 200 nm nanospheres reduced for 10, 20, ad 30 secs **(Figure S3, Supporting information)**. The samples used to evaluate the nanosphere size reduction presented in Figure 4 were coated with 24 nm of Al by thermal evaporation and the nanospheres were removed as discussed in the experimental details. The samples obtained were imaged in an SEM and the micrographs are shown in Figure 4c. Reducing the size of the nanospheres also minimizes defects such as merged pores which are critical for ensuring a narrow pore size in membranes. An example of this has been included in the supporting info **(Figure S3, Supporting information)**.

**Figure 4.**
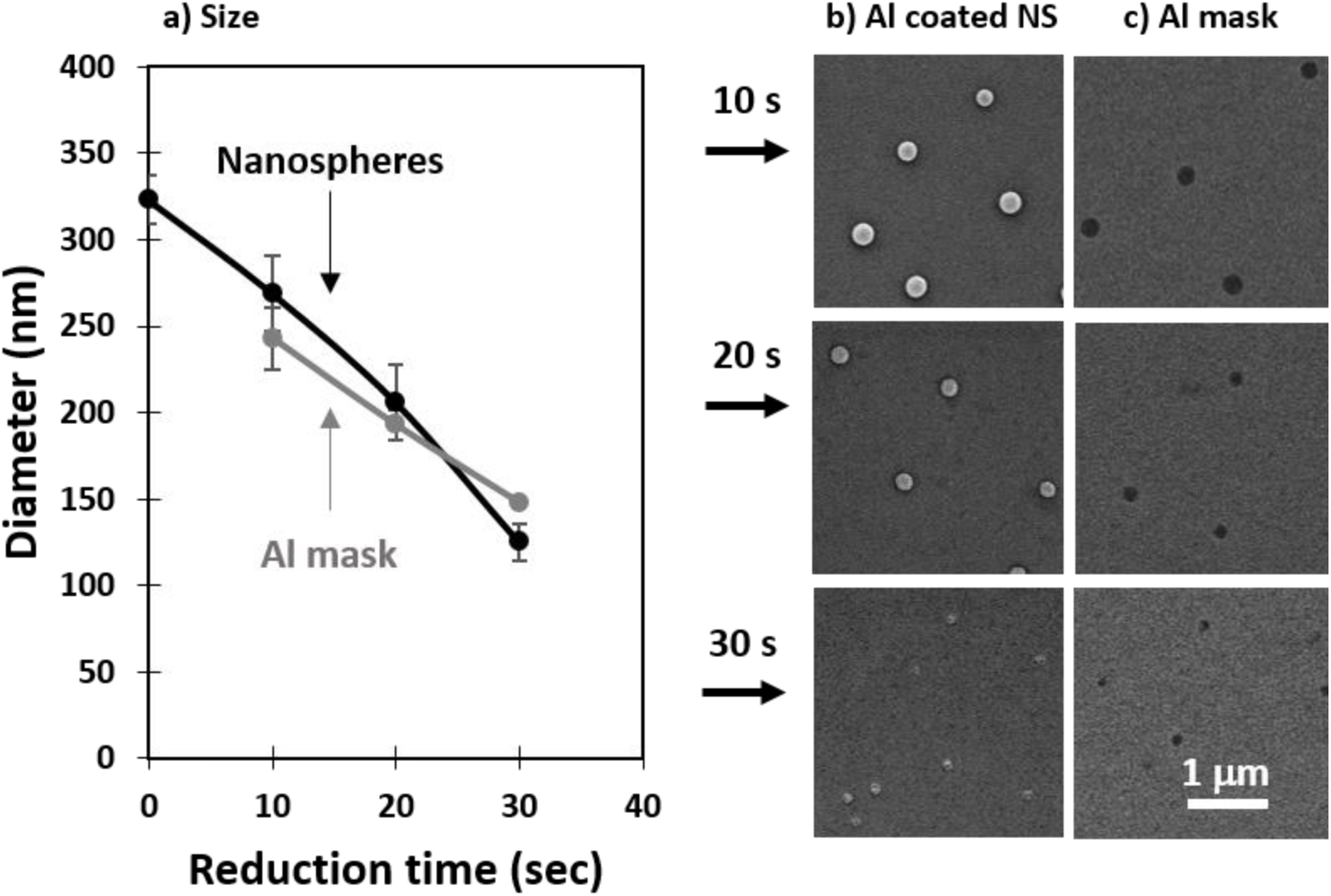
Nanosphere size reduction: a) diameter of the reduced nanospheres and the resulting nanopores in the Al mask, b) top-view of the reduced nanospheres, and c) the resulting pores in the Al mask.

In this work, we transferred the porous pattern produced in the Al masks from nanospheres reduced in size for 10, 20, and 30 s. We will now refer to these samples as membranes with small, medium, and large pores, respectively. The nanoporous pattern was transferred onto the underlying SiO_2_ film through RIE in an Ar:SF_6_ plasma using an adapted version of a previously reported recipe.^38,54^ After transferring the pattern, the Al was removed by wet etching in NaOH and the samples were imaged in an SEM. The shape of the nanospheres transfers to the pore geometry as seen in the top-view micrographs presented as insets in Figure 5a. As the nanospheres decrease in size and shrink they lose circularity and this can be seen transferred onto the smallest pores. The pore sizes were evaluated from the SEM images taken at a 10,000X magnification from 5 different areas and the results have been summarized as a histogram presented in Figure 5a. A comparison between the size of the reduced nanospheres and the resulting pores is presented in Figure 5c. The average measured pore diameters are in good agreement with the size of the nanospheres, deviating slightly for the largest pore size which is mainly due to the isotropic nature of the etching process.

**Figure 5.**
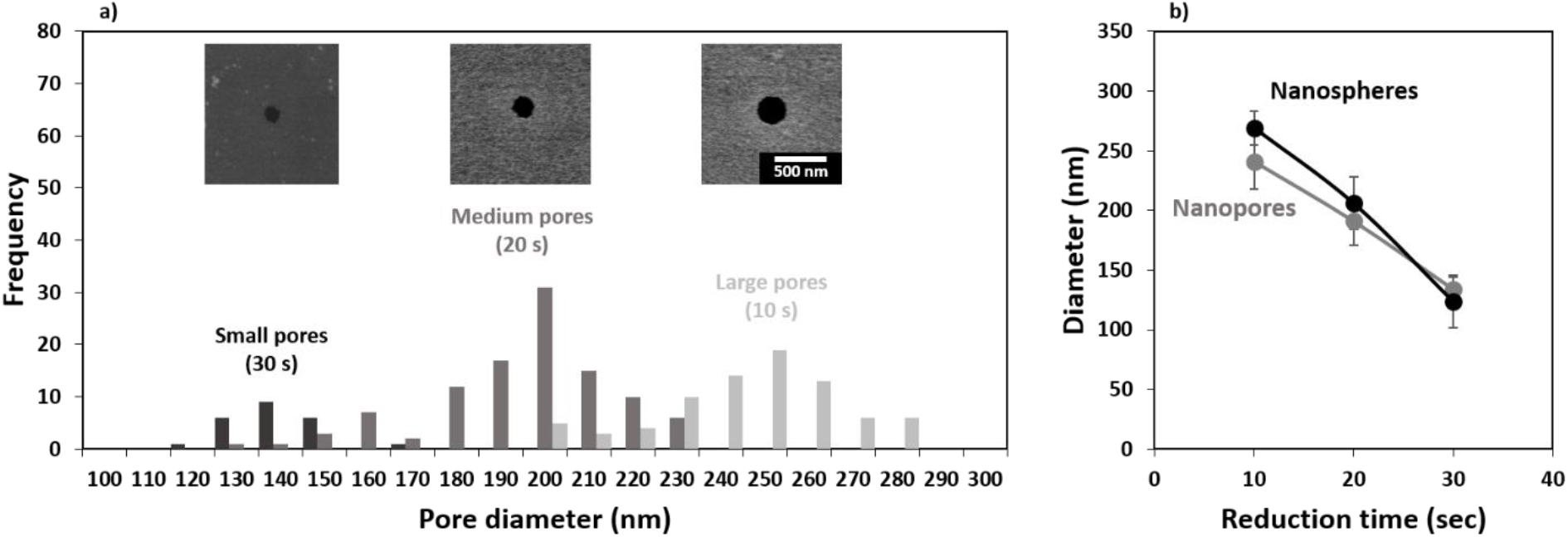
Final pore size as a function of nanosphere reduction time: a) data presented as a histogram from the pore sizes calculated from electronic imaging, b) representative SEM micrographs of the pores resulting from different nanosphere reduction times, and c) comparison between the reduced nanosphere size and the obtained pore size.

### 3.3 Size cut-off

The size cut-off of a membrane is a complex attribute, commonly measured experimentally for the species of interest. It has been shown that, in general, as the particle diameter approaches the size of a given circular pore the hindrance factor decreases from 1 to 0.^30,75^ A hindrance factor of 0 indicates full blockage of particles. One of the objectives in this work is to allow permeability of small EVs while limiting permeability of larger species (microvesicles and MVBs). We selected the membrane with 250 nm pores as these would show hindrance factors close to 1 for small EVs and close to 0 for large EVs. This membrane was integrated into a silicone-based device with top and bottom chambers separated by the membrane to evaluate its size cut-off. The devices were assembled following the process described in the experimental section (Figure 2). The top chamber was loaded with a testing solution containing either 200 or 300 nm fluorescent latex beads (unmixed). Figure 6a shows the size dispersion histogram obtained through nano particle tracking analysis (NTA) for the different beads used in the experiment. The particles with a nominal particle size of 200 nm showed an average measured diameter of 180 nm with a mode of 181 nm. The particles with a nominal particle size of 300 nm showed an average measured diameter of 296 nm with a mode of 290 nm. The testing species were allowed to permeate through the membrane into the bottom channel where a change in fluorescent intensity was tracked as function of time. The average intensity of the images has been plotted in Figure 6b, the values have been adjusted for background and plotted as the ratio I_F_/I_S_ which corresponds to the intensity recorded for the filtrate (I_F_) over the source (I_S_). I_F_ was measured in the bottom channel and I_S_ was measured at the center of the membrane. A size dispersion histogram of the pores in the membrane has also been included in Figure 6a. The 200 nm beads were able to pass through the membrane as evidenced by the increase in intensity seen in the bottom channel during the experiment. We did not observe an increase in fluorescent intensity in the bottom channel when testing the 300 nm beads. The latter is not unexpected and is due to the hindrance factor being zero or very close to zero for this case. For example, a 270 nm bead (smallest in the testing solution) and a 300 nm pore (largest in the membrane) show an approximate convective hindrance factor ≪ 0.05.^75^ This is in good agreement with the results shown in Figure 6b where the 300 nm beads have been blocked by the membrane.

**Figure 6.**
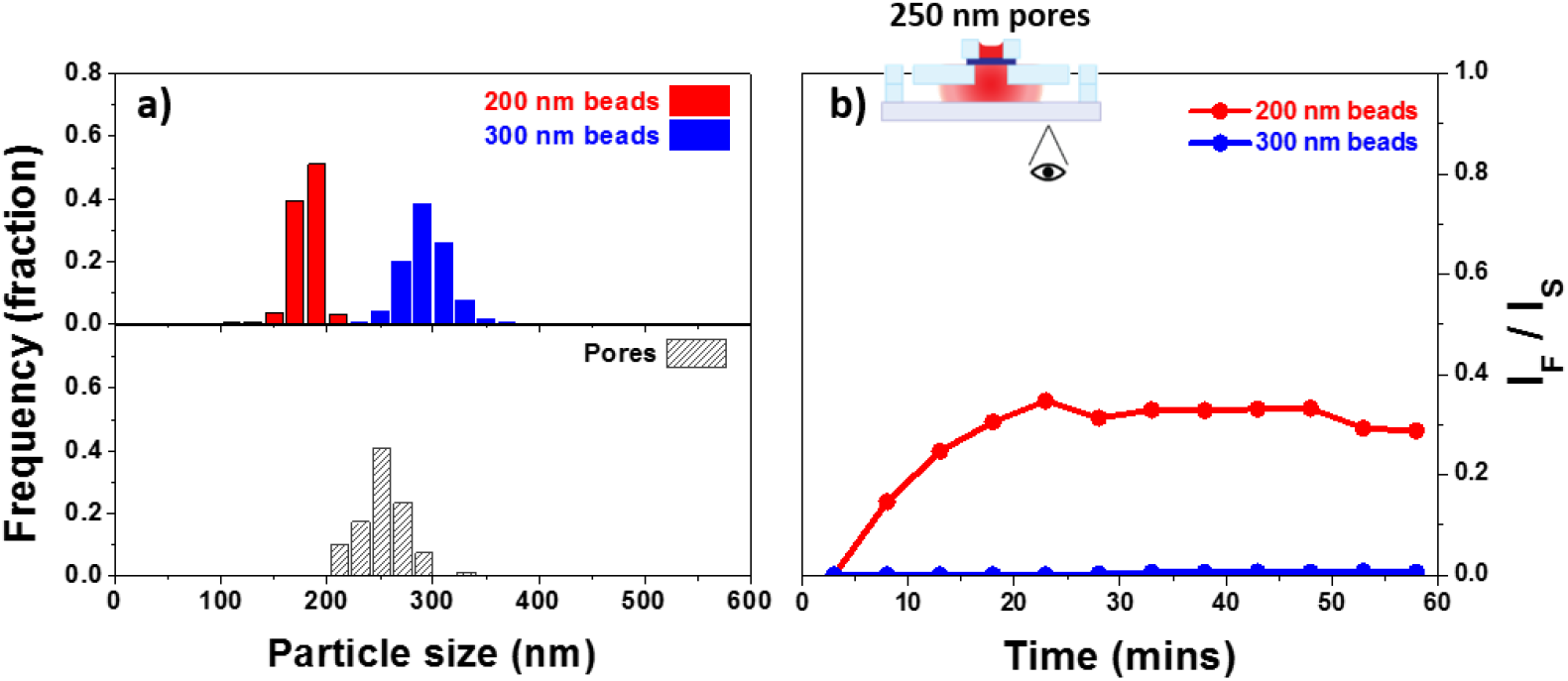
Size cut-off characteristics for a membrane of medium pore size-250 nm: a) size dispersion for the pores and the 200 and 300 nm beads used, and b) fluorescent intensity change tracked as I_F_/I_S_ for the permeating solution containing fluorescent beads of either size (unmixed).

### 3.4 Filtration of extracellular vesicles

Although polystyrene beads can be an acceptable model for EVs they are hard, rigid, and neutral in charge while EVs are soft, flexible, and exhibit surface charge. We further decided to evaluate the feasibility of utilizing our membranes for the study of EVs. As stated in the experimental details, this BSA coating was applied to neutralize the charges on the membrane therefore minimizing charge-based interaction. The EVs were fluorescently labelled with CFSE and allowed to permeate through a membrane with 250 nm pores coated with BSA. Figure 7a provides the size dispersion of the pores as well as the CFSE labelled EVs, source and filtrate, as measured by NTA. The fluorescent intensity in the bottom channel was tracked as function of time. The results have been background corrected and the ratio I_F_/I_S_ has been plotted in Figure 7b. The increase in intensity seen in Figure 7b indicated traffic across the membrane most likely due to small EVs, this was further confirmed using NTA. The size distribution curve for EVs narrows and shifts to smaller particle sizes after filtration. Quantitative determination of the NTA results was performed by comparing the mode and the mean of the particle size. The EVs source showed a mode value of 200 nm with a mean value of 253 nm. The filtered fraction recovered from the bottom channel showed a mode of 110 nm with a mean value of 166 nm which represents the small EVs. The membrane used here showed an effective size cut-off of 300 nm with the majority of the EVs ≤200 nm. These characteristics are in agreement with theoretical and experimental evidence of pores exhibiting an effective size cut-off smaller than their physical diameter.^30,75^ Moreover, because of minimal convective hindrance of EVs similar in size to the pores, EVs in the range between 200 and 300 nm represent a minor fraction. The observations found here demonstrate the capability of these membranes to modulate the permeability of EVs based on size.

**Figure 7.**
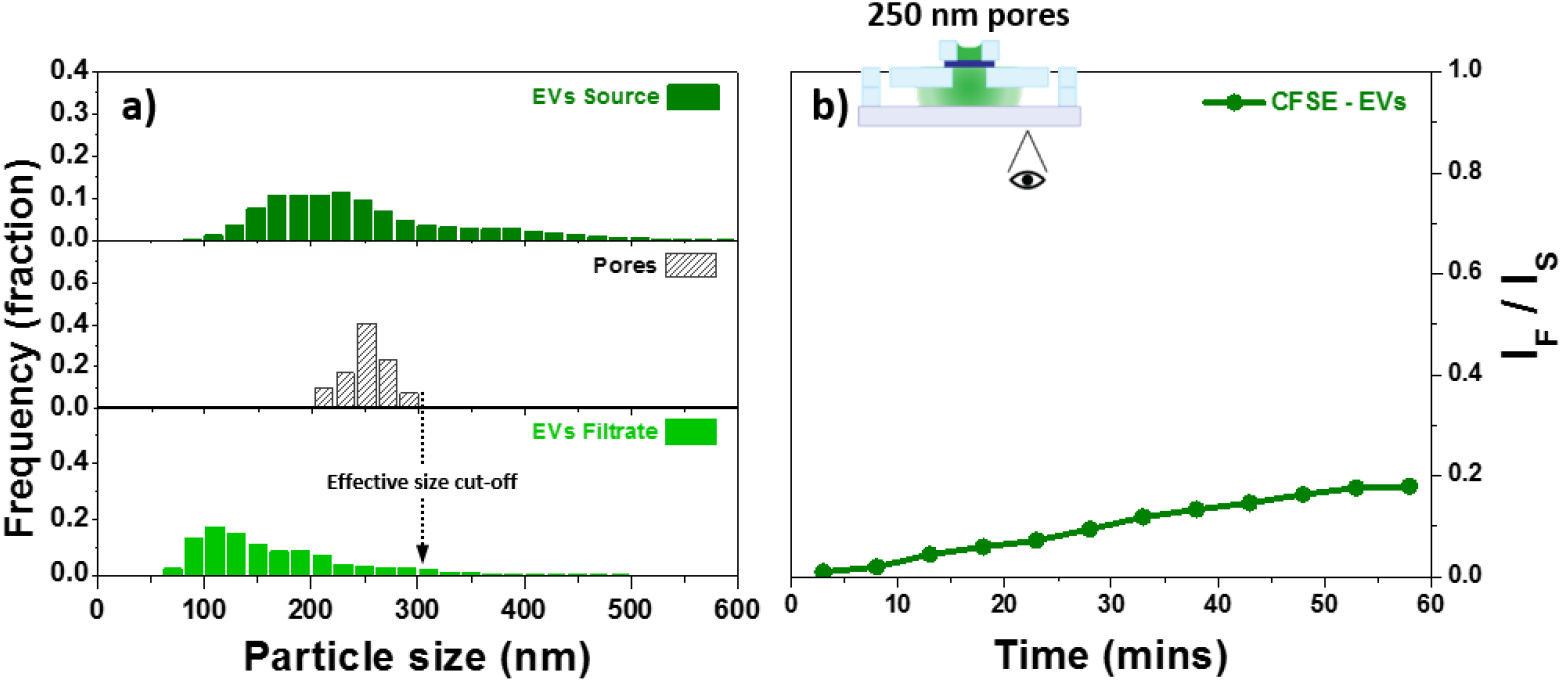
Filtration of EVs: a) size dispersion of the CFSE labelled EVs before and after filtration as measured by NTA, and b) fluorescent intensity increase on the bottom channel for EVs permeating through a membrane with 250 nm pores.

The evidence of the important role that EVs play in mediating cellular communication is undeniable.^76^ However, the vast majority of studies published to date have utilized polydisperse populations of EVs. A more sophisticated approach has become necessary, one that allows side-by-side comparison of the effect of different subtypes of EVs.^19^ The platform presented here fulfills this present need by offering an extensive process for the fabrication of ultrathin nanoporous membranes with the ability for pore size control over the full range of EVs.

Two attributes make this platform superior. Control over pore size, which determines the selectivity of the membrane to allow or block different subpopulations of EVs. Nanoscale thickness, which increases the permeability of species and allows for physical contact between cells across the membrane. These two characteristics are crucial for the improvement of current and emerging isolation and cellular co-culture platforms.^1,77,78^

Additionally, by simply changing the number of nanospheres introduced at the hexane-water interface one can control the overall spacing between pores. This is an advantageous attribute since it has been shown that membrane pore spacing can influence cell-substrate and cell-cell interactions.^79^

Lastly, the affordable and efficient nanoscale patterning and membrane lift-off methodologies utilized here also represent an additional improvement over current processes.

## 4 Conclusions

Here, we presented a process for the fabrication and integration of ultrathin nanoporous membranes with great potential for the study of EVs. At its basis, this process allows for pore size control into the nanoscale and offers a simple approach for membrane release. The membranes presented here achieved pore sizes in the range of small EVs and demonstrated their potential for the separation of EVs based on their size. This selectivity is crucial for filter-based isolation as well as for the modulation of EVs exchange in a co-culture systems. The ability to establish these models enables sophisticated interrogation of cellular behavior. Overall, this work provides a platform for the direct and indirect study of EVs for which interest will continue to grow.

## ASSOCIATED CONTENT

### Supporting information

Electron micrograph of SiO_2_ membranes fabricated with the same methodology are provided; as well as withdrawal speed optimization and nanosphere size reduction for 200 nm polystyrene nanospheres. Supporting information also includes a short video taken during membrane lift-off.

## ACKNOWLEDGEMENTS

Research reported in this publication was supported in part by NIGMS of the National Institutes of Health under award no. R35GM119623 to TRG.

## References

(1) Tkach, M.; Théry, C. Communication by Extracellular Vesicles: Where We Are and Where We Need to Go. Cell 2016, 164, 1226–1232.

(2) Ortiz, A. Not All Extracellular Vesicles Were Created Equal: Clinical Implications. Ann. Transl. Med. 2017, 5, 111–115.

(3) Colombo, M.; Raposo, G.; Théry, C. Biogenesis, Secretion, and Intercellular Interactions of Exosomes and Other Extracellular Vesicles. Annu. Rev. Cell Dev. Biol. 2014, 30, 255–289.

(4) Wang, W.; Luo, J.; Wang, S. Recent Progress in Isolation and Detection of Extracellular Vesicles for Cancer Diagnostics. Adv. Healthc. Mater. 2018, 7, 1800484–1800511.

(5) Hu, Y.; Li, D.; Wu, A.; Qiu, X.; Di, W.; Huang, L.; Qiu, L. TWEAK-Stimulated Macrophages Inhibit Metastasis of Epithelial Ovarian Cancer via Exosomal Shuttling of MicroRNA. Cancer Lett. 2017, 393, 60–67.

(6) Lucchetti, D.; Fattorossi, A.; Sgambato, A. Extracellular Vesicles in Oncology: Progress and Pitfalls in the Methods of Isolation and Analysis. Biotechnol. J. 2019, 14, 1–10.

(7) Kurywchak, P.; Tavormina, J.; Kalluri, R. The Emerging Roles of Exosomes in the Modulation of Immune Responses in Cancer. Genome Med. 2018, 10, 1–4.

(8) Peinado, H.; Aleckovic, M.; Lavotshkin, S.; Matei, I.; Costa-Silva, B.; Moreno-Bueno, G.; Hergueta-Redondo, M.; Williams, C.; Garcia-Santos, G.; Nitadori-Hoshino, A.; et al. Melanoma Exosomes Educate Bone Marrow Progenitor Cells toward a Pro-Metastatic Phenotype through MET. Nat. Med. 2013, 18, 883–891.

(9) Zhu, W.; Huang, L.; Li, Y.; Zhang, X.; Gu, J.; Yan, Y.; Xu, X.; Wang, M.; Qian, H.; Xu, W. Exosomes Derived from Human Bone Marrow Mesenchymal Stem Cells Promote Tumor Growth in Vivo. Cancer Lett. 2012, 315, 28–37.

(10) Wu, S.; Ju, G. Q.; Du, T.; Zhu, Y. J.; Liu, G. H. Microvesicles Derived from Human Umbilical Cord Wharton’s Jelly Mesenchymal Stem Cells Attenuate Bladder Tumor Cell Growth In Vitro and In Vivo. PLoS One 2013, 8, 1–12.

(11) Ono, M.; Kosaka, N.; Tominaga, N.; Yoshioka, Y.; Takeshita, F.; Takahashi, R.; Yoshida, M.; Tsuda, H.; Tamura, K.; Ochiya, T. Exosomes from Bone Marrow Mesenchymal Stem Cells Contain a MicroRNA That Promotes Dormancy in Metastatic Breast Cancer Cells. Sci. Signal. 2014, 7, ra63–ra63.

(12) Fonsato, V.; Collino, F.; Herrera, M. B.; Cavallari, C.; Deregibus, M. C.; Cisterna, B.; Bruno, S.; Romagnoli, R.; Salizzoni, M.; Tetta, C.; et al. Human Liver Stem Cell-Derived Microvesicles Inhibit Hepatoma Growth in SCID Mice by Delivering Antitumor MicroRNAs. Stem Cells 2012, 30, 1985–1998.

(13) Sun, W.; Zhao, C.; Li, Y.; Wang, L.; Nie, G.; Peng, J.; Wang, A.; Zhang, P.; Tian, W.; Li, Q.; et al. Osteoclast-Derived MicroRNA-Containing Exosomes Selectively Inhibit Osteoblast Activity. Cell Discov. 2016, 2, 1–23.

(14) Peak, T. C.; Praharaj, P. P.; Panigrahi, G. K.; Doyle, M.; Su, Y.; Schlaepfer, I. R.; Singh, R.; Vander Griend, D. J.; Alickson, J.; Hemal, A.; et al. Exosomes Secreted by Placental Stem Cells Selectively Inhibit Growth of Aggressive Prostate Cancer Cells. Biochem. Biophys. Res. Commun. 2018, 499, 1004–1010.

(15) Kalani, A.; Tyagi, A.; Tyagi, N. Exosomes: Mediators of Neurodegeneration, Neuroprotection and Therapeutics. Mol. Neurobiol. 2014, 49, 590–600.

(16) Van Niel, G.; D’Angelo, G.; Raposo, G. Shedding Light on the Cell Biology of Extracellular Vesicles. Nat. Rev. Mol. Cell Biol. 2018, 19, 213–228.

(17) Rani, S.; Ryan, A. E.; Griffin, M. D.; Ritter, T. Mesenchymal Stem Cell-Derived Extracellular Vesicles: Toward Cell-Free Therapeutic Applications. Mol. Ther. 2015, 23, 812–823.

(18) Witwer, K. W.; Van Balkom, B. W. M.; Bruno, S.; Choo, A.; Dominici, M.; Gimona, M.; Hill, A. F.; De Kleijn, D.; Koh, M.; Lai, R. C.; et al. Defining Mesenchymal Stromal Cell (MSC)-Derived Small Extracellular Vesicles for Therapeutic Applications. J. Extracell. Vesicles 2019, 8, 1609206.

(19) Tkach, M.; Kowal, J.; Théry, C. Why the Need and How to Approach the Functional Diversity of Extracellular Vesicles. Philos. Trans. R. Soc. B Biol. Sci. 2018, 373, 20160479.

(20) Shao, H.; Im, H.; Castro, C. M.; Breakefield, X.; Weissleder, R.; Lee, H. New Technologies for Analysis of Extracellular Vesicles. Chem. Rev. 2018, 118, 1917–1950.

(21) Hartjes, T. A.; Mytnyk, S.; Jenster, G. W.; van Steijn, V.; van Royen, M. E. Extracellular Vesicle Quantification and Characterization: Common Methods and Emerging Approaches. Bioengineering 2019, 6.

(22) Woo, H. K.; Sunkara, V.; Park, J.; Kim, T. H.; Han, J. R.; Kim, C. J.; Choi, H. Il; Kim, Y. K.; Cho, Y. K. Exodisc for Rapid, Size-Selective, and Efficient Isolation and Analysis of Nanoscale Extracellular Vesicles from Biological Samples. ACS Nano 2017, 11, 1360–1370.

(23) Liu, F.; Vermesh, O.; Mani, V.; Ge, T. J.; Madsen, S. J.; Sabour, A.; Hsu, E. C.; Gowrishankar, G.; Kanada, M.; Jokerst, J. V.; et al. The Exosome Total Isolation Chip. ACS Nano 2017, 11, 10712–10723.

(24) Heinemann, M. L.; Ilmer, M.; Silva, L. P.; Hawke, D. H.; Recio, A.; Vorontsova, M. A.; Alt, E.; Vykoukal, J. Benchtop Isolation and Characterization of Functional Exosomes by Sequential Filtration. J. Chromatogr. A 2014, 1371, 125–135.

(25) Gardiner, C.; Vizio, D. Di; Sahoo, S.; Théry, C.; Witwer, K. W.; Wauben, M.; Hill, A. F. Techniques Used for the Isolation and Characterization of Extracellular Vesicles: Results of a Worldwide Survey. J. Extracell. Vesicles 2016, 5, 32945.

(26) Konoshenko, M. Y.; Lekchnov, E. A.; Vlassov, A. V.; Laktionov, P. P. Isolation of Extracellular Vesicles: General Methodologies and Latest Trends. Biomed Res. Int. 2018, 2018.

(27) Lu, J.; Pang, J.; Chen, Y.; Dong, Q.; Sheng, J.; Luo, Y.; Lu, Y.; Lin, B.; Liu, T. Application of Microfluidic Chips in Separation and Analysis of Extracellular Vesicles in Liquid Biopsy for Cancer. Micromachines 2019, 10, 390.

(28) Guo, S. C.; Tao, S. C.; Dawn, H. Microfluidics-Based on-a-Chip Systems for Isolating and Analysing Extracellular Vesicles. J. Extracell. Vesicles 2018, 7, 1508271.

(29) Takov, K.; Yellon, D. M.; Davidson, S. M. Comparison of Small Extracellular Vesicles Isolated from Plasma by Ultracentrifugation or Size-Exclusion Chromatography: Yield, Purity and Functional Potential. J. Extracell. Vesicles 2019, 8, 1560809.

(30) Snyder, J. L.; Clark, A.; Fang, D. Z.; Gaborski, T. R.; Striemer, C. C.; Fauchet, P. M.; McGrath, J. L. An Experimental and Theoretical Analysis of Molecular Separations by Diffusion through Ultrathin Nanoporous Membranes. J. Memb. Sci. 2011, 369, 119–129.

(31) Mehta, A.; Zydney, A. L. Permeability and Selectivity Analysis for Ultrafiltration Membranes. J. Memb. Sci. 2005, 249, 245–249.

(32) Apel, P. Y.; Blonskaya, I. V; Dmitriev, S. N.; Orelovitch, O. L.; Sartowska, B. Structure of Polycarbonate Track-Etch Membranes: Origin of the “Paradoxical” Pore Shape. J. Memb. Sci. 2006, 282, 393–400.

(33) Gaborski, T. R.; Snyder, J. L.; Striemer, C. C.; Fang, D. Z.; Hoffman, M.; Fauchet, P. M.; McGrath, J. L. High-Performance Separation of Nanoparticles with Ultrathin Porous Nanocrystalline Silicon Membranes. ACS Nano 2010, 4, 6973–6981.

(34) Chung, H. H.; Mireles, M.; Kwarta, B. J.; Gaborski, T. R. Use of Porous Membranes in Tissue Barrier and Co-Culture Models. Lab Chip 2018, 18, 1671–1689.

(35) Mireles, M.; Gaborski, T. R. Fabrication Techniques Enabling Ultrathin Nanostructured Membranes for Separations. Electrophoresis 2017, 38, 2374–2388.

(36) Mazzocchi, A. R.; Man, A. J.; DesOrmeaux, J.-P. S.; Gaborski, T. R. Porous Membranes Promote Endothelial Differentiation of Adipose-Derived Stem Cells and Perivascular Interactions. Cell. Mol. Bioeng. 2014, 7, 369–378.

(37) Ryu, S.; Yoo, J.; Jang, Y.; Han, J.; Yu, S. J.; Park, J.; Jung, S. Y.; Ahn, K. H.; Im, S. G.; Char, K.; et al. Nanothin Coculture Membranes with Tunable Pore Architecture and Thermoresponsive Functionality for Transfer-Printable Stem Cell-Derived Cardiac Sheets. ACS Nano 2015, 9, 10186–10202.

(38) Carter, R. N.; Casillo, S. M.; Mazzocchi, A. R.; DesOrmeaux, J.-P. S.; Roussie, J. A.; Gaborski, T. R. Ultrathin Transparent Membranes for Cellular Barrier and Co-Culture Models. Biofabrication 2017, 9, 15019.

(39) Sainiemi, L.; Viheriälä, J.; Sikanen, T.; Laukkanen, J.; Niemi, T. Nanoperforated Silicon Membranes Fabricated by UV-Nanoimprint Lithography, Deep Reactive Ion Etching and Atomic Layer Deposition. J. Micromechanics Microengineering 2010, 20, 077001.

(40) Striemer, C. C.; Gaborski, T. R.; McGrath, J. L.; Fauchet, P. M. Charge-and Size-Based Separation of Macromolecules Using Ultrathin Silicon Membranes. Nature 2007, 445, 749–753.

(41) Brassat, K.; Kool, D. Hierarchical Nanopores Formed by Block Copolymer Lithography on the Surfaces of Different Materials Pre-Patterned by Nanosphere Lithography. Nanoscale 2018, 10, 10005–10017.

(42) Wong, H. C.; Zhang, Y.; Viasnoff, V.; Low, H. Y. Predictive Design, Etch-Free Fabrication of Through-Hole Membrane with Ordered Pores and Hierarchical Layer Structure. Adv. Mater. Technol. 2017, 2, 1600169.

(43) Wong, H. C.; Grenci, G.; Wu, J.; Viasnoff, V.; Low, H. Y. Roll-to-Roll Fabrication of Residual-Layer-Free Micro/Nanoscale Membranes with Precise Pore Architectures and Tunable Surface Textures. Ind. Eng. Chem. Res. 2018, 57, 13759–13768.

(44) Nabar, B. P.; Çelik-Butler, Z.; Dennis, B. H.; Billo, R. E. A Nanoporous Silicon Nitride Membrane Using a Two-Step Lift-off Pattern Transfer with Thermal Nanoimprint Lithography. J. Micromechanics Microengineering 2012, 22, 45012.

(45) Montagne, F.; Blondiaux, N.; Bojko, A.; Pugin, R. Molecular Transport through Nanoporous Silicon Nitride Membranes Produced from Self-Assembling Block Copolymers. Nanoscale 2012, 4, 5880–5886.

(46) Acikgoz, C.; Ling, X. Y.; Phang, I. Y.; Hempenius, M. A.; Reinhoudt, D. N.; Huskens, J.; Vancso, G. J. Fabrication of Freestanding Nanoporous Polyethersulfone Membranes Using Organometallic Polymer Resists Patterned by Nanosphere Lithography. Adv. Mater. 2009, 21, 2064–2067.

(47) Nuxoll, E. E.; Hillmyer, M. A.; Wang, R.; Leighton, C.; Siegel, R. A. Composite Block Polymer-Microfabricated Silicon Nanoporous Membrane. ACS Appl. Mater. Interfaces 2009, 1, 888–893.

(48) Wang, Y.; Li, F. An Emerging Pore-Making Strategy: Confined Swelling-Induced Pore Generation in Block Copolymer Materials. Adv. Mater. 2011, 23, 2134–2148.

(49) Yu, H.; Qiu, X.; Nunes, S. P.; Peinemann, K. V. Self-Assembled Isoporous Block Copolymer Membranes with Tuned Pore Sizes. Angew. Chemie - Int. Ed. 2014, 53, 10072–10076.

(50) Altinpinar, S.; Zhao, H.; Ali, W.; Kappes, R. S.; Schuchardt, P.; Salehi, S.; Santoro, G.; Theato, P.; Roth, S. V.; Gutmann, J. S. Distortion of Ultrathin Photocleavable Block Copolymer Films during Photocleavage and Nanopore Formation. Langmuir 2015, 31, 8947–8952.

(51) Isa, L.; Kumar, K.; Müller, M.; Grolig, J.; Textor, M.; Reimhult, E. Particle Lithography from Colloidal Self-Assembly at Liquid-Liquid Interfaces. ACS Nano 2010, 4, 5665–5670.

(52) Kang, C.; Ramakrishna, S. N.; Nelson, A.; Cremmel, C. V. M.; vom Stein, H.; Spencer, N. D.; Isa, L.; Benetti, E. M. Ultrathin, Freestanding, Stimuli-Responsive, Porous Membranes from Polymer Hydrogel-Brushes. Nanoscale 2015, 7, 13017–13025.

(53) Klein, M. J. K.; Montagne, F.; Blondiaux, N.; Vazquez-Mena, O.; Heinzelmann, H.; Pugin, R.; Brugger, J.; Savu, V. SiN Membranes with Submicrometer Hole Arrays Patterned by Wafer-Scale Nanosphere Lithographya) SiN Membrane Masks for X‐ray Lithography SiN Membranes with Submicrometer Hole Arrays Patterned by Wafer-Scale Nanosphere Lithography. J. Vac. Sci. {&} Technol. B J. Appl. Phys. Phys. Lett. J. Vac. Sci. Technol 2011, 29, 21012–22599.

(54) Miller, J. J.; Carter, R. N.; McNabb, K. B.; DesOrmeaux, J.-P. S.; Striemer, C. C.; Winans, J. D.; Gaborski, T. R. Lift-off of Large-Scale Ultrathin Nanomembranes. J. Micromechanics Microengineering 2015, 25, 015011.

(55) Dehghani, M.; Lucas, K.; Flax, J.; McGrath, J.; Gaborski, T. Tangential Flow Microfluidics for the Capture and Release of Nanoparticles and Extracellular Vesicles on Conventional and Ultrathin Membranes. bioRxiv 2019.

(56) Lotito, V.; Zambelli, T. Approaches to Self-Assembly of Colloidal Monolayers: A Guide for Nanotechnologists. Adv. Colloid Interface Sci. 2017, 246, 217–274.

(57) Sang Han, G.; Lee, S.; Un Jin, Y.; Sun Cho, I.; Suk Jung, H. Facile Transfer Fabrication of Transparent, Conductive and Flexible In2O3:Sn (ITO) Nanowire Arrays Electrode via Selective Wet-Etching ZnO Sacrificial Layer. Mater. Lett. 2015, 158, 304–308.

(58) Rajan, A.; Rogers, D. J.; Ton-That, C.; Zhu, L.; Phillips, M. R.; Sundaram, S.; Gautier, S.; Moudakir, T.; El-Gmili, Y.; Ougazzaden, A.; et al. Wafer-Scale Epitaxial Lift-off of Optoelectronic Grade GaN from a GaN Substrate Using a Sacrificial ZnO Interlayer. J. Phys. D. Appl. Phys. 2016, 49, 315105.

(59) Rogers, D. J.; Ougazzaden, A.; Sandana, V. E.; Moudakir, T.; Ahaitouf, A.; Teherani, F. H.; Gautier, S.; Goubert, L.; Davidson, I. a.; Prior, K. a.; et al. Novel Process for Direct Bonding of GaN onto Glass Substrates Using Sacrificial ZnO Template Layers to Chemically Lift-off GaN from c-Sapphire. In Oxide-based Materials and Devices III. International Society for Optics and Photonics.; 2012; Vol. 8263, p 82630R–1.

(60) Kim, M. Y.; Li, D. J.; Pham, L. K.; Wong, B. G.; Hui, E. E. Microfabrication of High-Resolution Porous Membranes for Cell Culture. J. Memb. Sci. 2014, 452, 460–469.

(61) Elnathan, R.; Isa, L.; Brodoceanu, D.; Nelson, A.; Harding, F. J.; Delalat, B.; Kraus, T.; Voelcker, N. H. Versatile Particle-Based Route to Engineer Vertically Aligned Silicon Nanowire Arrays and Nanoscale Pores. ACS Appl. Mater. Interfaces 2015, 7, 23717–23724.

(62) Rey, B. M.; Elnathan, R.; Ditcovski, R.; Geisel, K.; Zanini, M.; Fernandez-Rodriguez, M. A.; Naik, V. V.; Frutiger, A.; Richtering, W.; Ellenbogen, T.; et al. Fully Tunable Silicon Nanowire Arrays Fabricated by Soft Nanoparticle Templating. Nano Lett. 2016, 16, 157–163.

(63) Morales-Kastresana, A.; Telford, B.; Musich, T. A.; McKinnon, K.; Clayborne, C.; Braig, Z.; Rosner, A.; Demberg, T.; Watson, D. C.; Karpova, T. S.; et al. Labeling Extracellular Vesicles for Nanoscale Flow Cytometry. Sci. Rep. 2017, 7, 1–10.

(64) Quah, B. J. C.; Parish, C. R. The Use of Carboxyfluorescein Diacetate Succinimidyl Ester (CFSE) to Monitor Lymphocyte Proliferation. J. Vis. Exp. 2010, 44, e2259.

(65) Liu, D.; Yu, J.; Chen, H.; Reichman, R.; Wu, H.; Jin, X. Statistical Determination of Threshold for Cellular Division in the CFSE-Labeling Assay. J. Immunol. Methods 2006, 312, 126–136.

(66) Parish, C. R.; Glidden, M. H.; Quah, B. J. C.; Warren, H. S. Use of the Intracellular Fluorescent Dye CFSE to Monitor Lymphocyte Migration and Proliferatio. Curr. Protoc. Immunol. 2009, 84, 4–9.

(67) Longatti, A.; Schindler, C.; Collinson, A.; Jenkinson, L.; Matthews, C.; Fitzpatrick, L.; Blundy, M.; Minter, R.; Vaughan, T.; Shaw, M.; et al. High Affinity Single-Chain Variable Fragments Are Specific and Versatile Targeting Motifs for Extracellular Vesicles. Nanoscale 2018, 10, 14230–14244.

(68) Keller, S.; König, A. K.; Marmé, F.; Runz, S.; Wolterink, S.; Koensgen, D.; Mustea, A.; Sehouli, J.; Altevogt, P. Systemic Presence and Tumor-Growth Promoting Effect of Ovarian Carcinoma Released Exosomes. Cancer Lett. 2009, 278, 73–81.

(69) Pospichalova, V.; Svoboda, J.; Dave, Z.; Kotrbova, A.; Kaiser, K.; Klemova, D.; Ilkovics, L.; Hampl, A.; Crha, I.; Jandakova, E.; et al. Simplified Protocol for Flow Cytometry Analysis of Fluorescently Labeled Exosomes and Microvesicles Using Dedicated Flow Cytometer. J. Extracell. Vesicles 2015, 4, 1–15.

(70) Dehghani, M.; Gulvin, S. M.; Flax, J.; Thomas, R. Exosome Labeling by Lipophilic Dye PKH26 Results in Significant Increase in Vesicle Size. BioRxiv 2019.

(71) Vogel, N.; Retsch, M.; Fustin, C. A.; Del Campo, A.; Jonas, U. Advances in Colloidal Assembly: The Design of Structure and Hierarchy in Two and Three Dimensions. Chem. Rev. 2015, 115, 6265–6311.

(72) Blättler, T. M.; Binkert, A.; Zimmermann, M.; Textor, M.; Vörös, J.; Reimhult, E. From Particle Self-Assembly to Functionalized Sub-Micron Protein Patterns. Nanotechnology 2008, 19, 075301.

(73) Plettl, A.; Enderle, F.; Saitner, M.; Manzke, A.; Pfahler, C.; Wiedemann, S.; Ziemann, P. Non-Close-Packed Crystals from Self-Assembled Polystyrene Spheres by Isotropic Plasma Etching: Adding Flexibility to Colloid Lithography. Adv. Funct. Mater. 2009, 19, 3279–3284.

(74) Vogel, N.; Goerres, S.; Landfester, K.; Weiss, C. K. A Convenient Method to Produce Close- and Non-Close-Packed Monolayers Using Direct Assembly at the Air-Water Interface and Subsequent Plasma-Induced Size Reduction. Macromol. Chem. Phys. 2011, 212, 1719–1734.

(75) Dechadilok, P.; Deen, W. M. Hindrance Factors for Diffusion and Convection in Pores. 2006, 6953–6959.

(76) Mathieu, M.; Martin-Jaular, L.; Lavieu, G.; Théry, C. Specificities of Secretion and Uptake of Exosomes and Other Extracellular Vesicles for Cell-to-Cell Communication. Nat. Cell Biol. 2019, 21, 9–17.

(77) Li, P.; Kaslan, M.; Lee, S. H.; Yao, J.; Gao, Z. Progress in Exosome Isolation Techniques. Theranostics 2017, 7, 789–804.

(78) Théry, C.; Amigorena, S.; Raposo, G.; Clayton, A. Isolation and Characterization of Exosomes from Cell Culture Supernatants and Biological Fluids. Curr. Protoc. Cell Biol. 2006, 30, 3.22.1–3.22.29.

(79) Casillo, S. M.; Peredo, A. P.; Perry, S. J.; Chung, H. H.; Gaborski, T. R. Membrane Pore Spacing Can Modulate Endothelial Cell–Substrate and Cell–Cell Interactions. ACS Biomater. Sci. Eng. 2017, 3, 243–248.

