## Supplemental Figures for "Use of Nanosphere Self-Assembly to Pattern Nanoporous Membranes for the Study of Extracellular Vesicles"

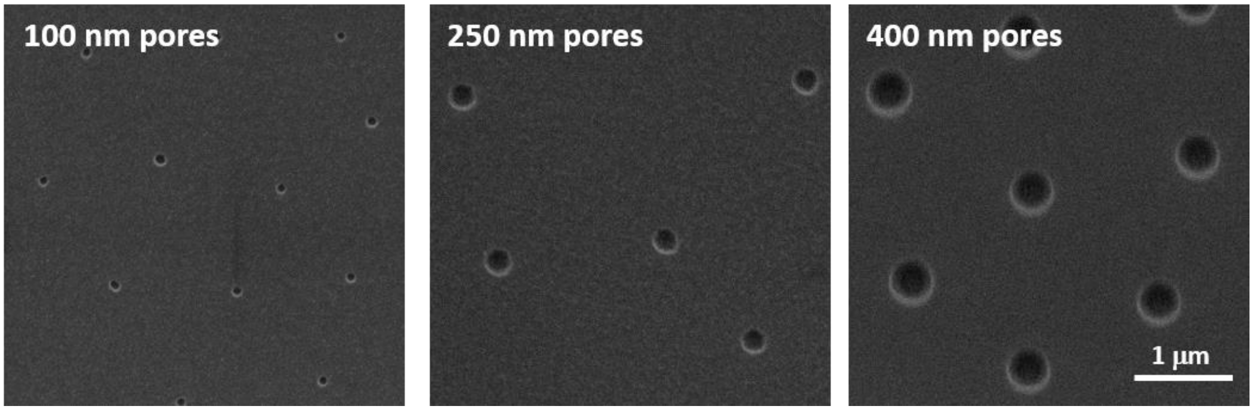


**Figure S1. $SiO_2$ membranes.** Top-vie electron micrographs taken from three different $SiO_2$ membranes fabricated following the same procedure described for SiN in the main manuscript.

In this case, the $SiO_2$ was deposited by plasma enhanced chemical vapor deposition in an Applied Materials P500 tool at 390 $^oC$ and a working pressure of 1.2 kPa using tetraethoxysilane (TEOS) as precursor. After $SiO_2$ deposition, the stack (Si/ZnO/$SiO_2$) was annealed at 600 $^oC$ under $N_2$ flow for 1 h for stress stabilization. This $SiO_2$ deposition and subsequent annealing has been previously found to yield a tensile film stress of 150 MPa, stable over time.[1]

(1) Carter, R. N.; Casillo, S. M.; Mazzocchi, A. R.; DesOrmeaux, J.-P. S.; Roussie, J. A.; Gaborski, T. R. Ultrathin Transparent Membranes for Cellular Barrier and Co-Culture Models. *Biofabrication* **2017**, *9*, 15019.

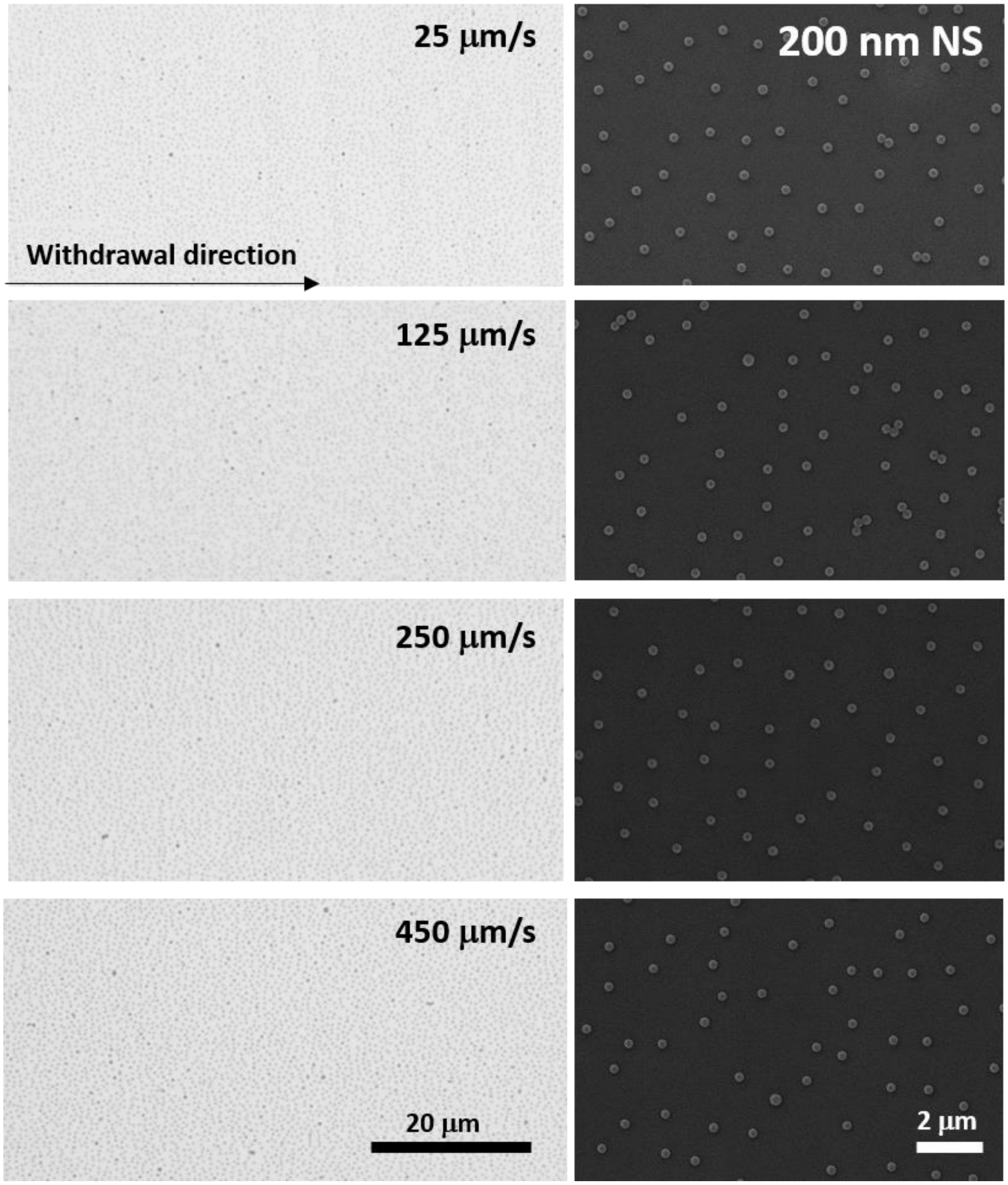


**Figure S2. Withdrawal speed optimization.** Optical and electron micrographs taken from 200 nm self-assembled monolayers transferred onto a substrate at different withdrawal speeds.

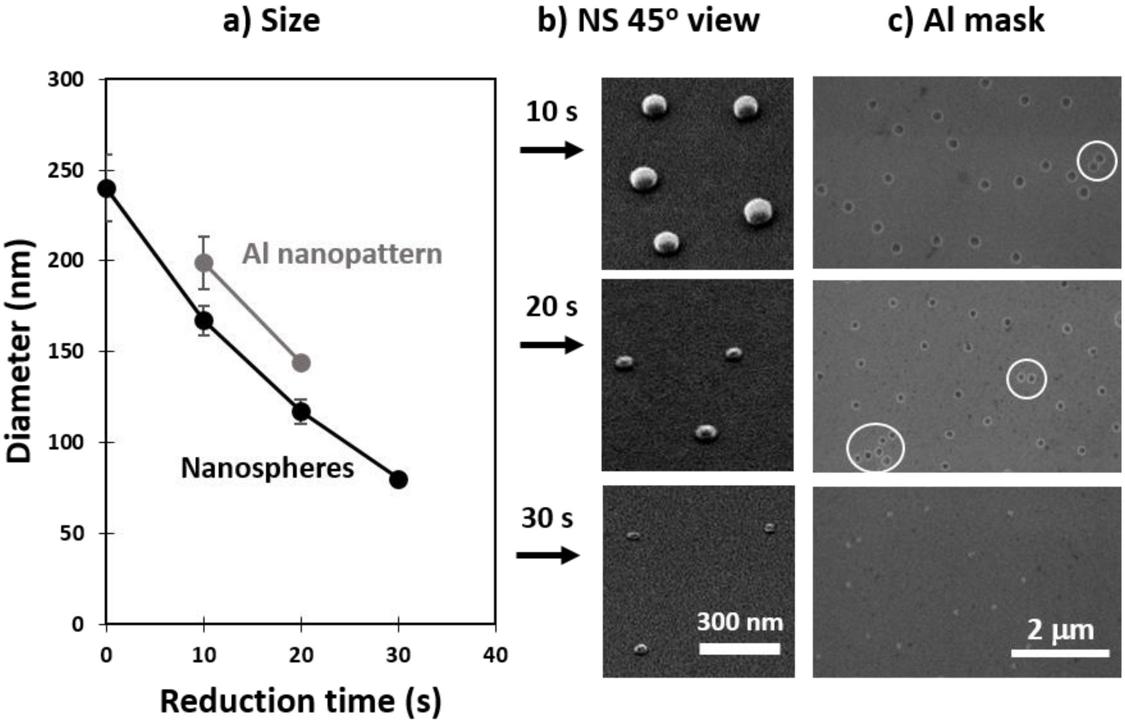


**Figure S3. Nanosphere size reduction:** a) diameter of the reduced nanospheres (200 nm nominal size) and the resulting nanopores in the Al mask, b) tilted-view of the reduced nanospheres, and c) the resulting pores in the Al mask and unmerging of pores.

The Al mask produced after 10 s of nanosphere reduction presents a doublet (circled in white). When reducing the nanospheres for 20 s two individual pores are observed in close proximity, these constitute a doublet that has been unmerged after reducing the size of the nanospheres.

**Supporting video 1. Membrane lift-off:** following immersion in 1M HCl for sacrificial film removal, an ultrathin nanoporous silico nitride membrane is released from its supporting substrate, Si wafer, with the use of tweezers.
